# Successful optimization of CRISPR/Cas9-mediated defined point mutation knock-in using allele-specific PCR assays in zebrafish

**DOI:** 10.1101/194936

**Authors:** Sergey V. Prykhozhij, Charlotte Fuller, Shelby L. Steele, Chansey J. Veinotte, Babak Razaghi, Johane Robitaille, Christopher McMaster, Adam Shlien, David Malkin, Jason N. Berman

## Abstract

Single-stranded oligodeoxynucleotides (ssODN) are donor templates for homology-directed repair-based knock-in of point mutations using CRISPR/Cas9. To optimize the efficiency of ssODN-based knock-ins in zebrafish, we developed allele-specific PCR (AS-PCR) assays for introducing point mutations in *tp53*, *cdh5* and *lmna* as case studies. In these point mutation strategies we introduced the codon mutations, sgRNA site mutations and restriction sites which can be detected by AS-PCR with the primers matching their respective alleles in combination with a common primer. We employed the anti-sense asymmetric oligo design as the main optimization as well as phosphorothioate oligo modification and also observed that proximity of the mutation site to the Cas9 cut site improves the efficiency when knock-ins into different genes were compared. We improved the efficiencies of two *tp53* knock-ins using anti-sense asymmetric ultramer oligos (126-nt in length with homology arms of 36 and 90 nucleotides, anti-sense to the sgRNA) by 3-10 fold, the optimizations which resulted in successful founders for both *tp53* knock-ins with transmission rates of 20-40 %. The initially low knock-in efficiency for *tp53* mutants was likely due to the distance between the Cas9 cut site and mutations since *cdh5* G767S knock-in located at the cut site had much higher founder identification and germline transmission rates. The phosphorothioate oligo modifications was used for a lamin A/C (*lmna*) knock-in strategy and it resulted in 40 % overall improvement in knock-in efficiency and greater knock-in consistency. We also determined that AS-PCR detected false-positive knock-ins which constituted 25-80 % of total in different strategies and developed a workflow to screen out the founders and F1 zebrafish carrying these undesirable modifications. In summary, we provide a complementary set of optimizations for CRISPR/Cas9-based ssODN knock-ins in zebrafish using a novel combination of methods.

## Introduction

Development and application of CRISPR/Cas9 technology has now been generally accepted as one of the most revolutionary biotechnologies of the last decade. In addition, to the fact that CRISPR/Cas9 can be used in a large variety of species, the most significant reason for its power is the ability to introduce specific modifications. However, introducing defined mutations has been a significant challenge in some species because it requires some form of homology-directed repair or recombination. Zebrafish (*Danio rerio*) has been one such species despite a wide adoption of CRISPR/Cas9 technology overall and significant advances in many areas^1–4^. Zebrafish is frequently used for basic research and disease modeling because of its transparency, fecundity, developed genetic and cell biological tools among other advantages. In terms of disease modeling, application of CRISPR/Cas9 to generate point missense mutants of residues conserved between humans and zebrafish can be particularly interesting as these types of studies in zebrafish can be significantly more cost-effective and scalable than in other vertebrate model animals, such as mice. However, this potential can only be realized if efficiencies of point mutation knock-in strategies can be substantially improved.

The first demonstrations of small mutation knock-ins in zebrafish^5,6^ were proof-of-concept that such modifications using ssODNs are possible, but did not show that the introduced mutations could actually be transmitted to the next generation. Importantly, the feasibility of introducing small modifications using ssODNs was established much earlier by previous groundbreaking work employing TALENs^7,8^. However, TALEN-based methods were not widely adopted by the field for knock-in generation, likely due to the greater difficulty of using TALENs than producing sgRNAs once CRISPR/Cas9 became available. Over the last few years, ssODN-based knock-ins advanced in several respects. Introduction of inserts encoding protein epitope tags such as HA was successfully accomplished, although the proportion of correctly-modified alleles was low^9,10^. This problem of low-fidelity insertions occurred in all early studies on ssODN-based knock-ins and no systematic attempt to address this problem has yet been published. On the other hand, the first report of point mutation insertion and germline transmission was published, describing the modification of *tardp* and *fus* genes involved in amyotrophic lateral sclerosis^11^. These authors were able to introduce a mutation into *fus* gene in 1 of 47 founders using a 33-nucleotide (nt) oligo and another mutation into *tardp* gene in 3 out of 77 founders using a 100-nt oligo containing sgRNA site mutations. None of the successful knock-ins contained indels but whether this is a representative sample from a real distribution of knock-in alleles is unclear. Next-generation sequencing methods will likely help establish how accurate different point mutation knock-in methods are. This paper by Armstrong et al. (2016)^11^ is important in that it shows that knock-ins are certainly doable and germline transmissible but its methodology is heavily reliant on sequencing of many PCR products without an easier prescreening method.

Several important optimizations emerged in recent publications on CRISPR/Cas9-mediated knock-ins that will likely impact similar work in zebrafish. The findings in two recent papers^12,13^ suggest the strong inverse relationship between the knock-in efficiency and the distance of the modification to the cut site. The most efficient positions for modification should be located < 15 nucleotides and ideally < 10 nucleotides away from the cut site as at a distance of 20 nucleotides away from the cut site, the efficiency drops to 20-30 % of the maximum as observed in the mouse cells^13^. Another important trend has been the focus on the structure of oligos. Asymmetric anti-sense oligos with homology arms of 36 and 90 nt were demonstrated by Richardson et al. (2016)^14^ to be superior to all other designs of the same size. The oligos in this case are anti-sense to the PAM-containing (non-target) strand, a portion of which was proposed to separate from the Cas9-sgRNA RNP and become available to bind the 36-nt homology arm of the oligo. This event can then facilitate homology-directed repair, as evidenced by the highly efficient repair of a mutated EGFP gene^14^. Chemical modifications have also been applied to ssODNs. Two phosphorothioate (phosphate where an oxygen is replaced with a sulphur atom) (PS) linkages at the ends of ssODNs promote generation of knock-ins in human cell lines compared to the oligos with traditional phosphate-oxygen bonds. Knock-in stimulation was regardless of the size of the oligos most likely due to blocking the activity of exonucleases^15^. This result was confirmed in other studies of knock-ins in cell lines^12,16,17^. PS-modification of oligos has also been tried in zebrafish^15^, but these authors found mainly imprecise knock-ins and did not seek direct evidence of knock-in stimulation. Thus, it is still unclear if PS modifications can be applied in zebrafish knock-in experiments.

In this manuscript, we apply these modifications to improve the ease and efficiency of performing point mutation knock-ins in zebrafish. For detection, we assessed the relative performance of restriction site-based measurement using sites introduced by knock-ins with synonymous mutation and allele-specific PCR (AS-PCR) assays most commonly used for nucleotide polymorphism detection^18^. We found that AS-PCR has a dramatically greater sensitivity than the restriction-based method. For achieving improved rates of knock-in generation, we took advantage of the proximity of the cut site to the site of modification in the case of *cdh5* G767S mutation, but for *tp53* mutations, this was not feasible. Therefore, we switched to asymmetric anti-sense oligos, which resulted in dramatic improvement (3-10 fold) of knock-in rates as measured by AS-PCR and next-generation sequencing. Upon isolation of zebrafish with correct knock-ins, we confirmed them at the genomic DNA and cDNA levels. This new method of knock-in genotyping also led us to identify an off-target phenomenon of *trans* knock-ins, when oligos used for knock-ins were inserted into other loci but could still generate false-positive AS-PCR hits. To facilitate the process of distinguishing true knock-ins from *trans* knock-ins, we used a combination of AS-PCR followed by restriction digestion of PCR amplicons centered on the knock-in sites at the restriction sites introduced as synonymous mutations. Finally, for the lamin A/C (*lmna*) knock-in strategies, we demonstrated that phosphorothioate oligos produce a significant improvement in knock-in efficiency and consistency as measured by AS-PCR. In sum, by applying these strategies, we have optimized CRISPR/Cas9-based knock ins in the zebrafish, enhancing the genome editing toolbox for zebrafish researchers to more efficiently model human genetic disorders.

## Materials and Methods

### Animal care and husbandry

Zebrafish housing, breeding conditions, and developmental staging of larvae, were performed according to Westerfield^19^. Use of zebrafish in this study was approved by the Dalhousie University Committee on Laboratory Animals (Protocols 15-125, 15-134). All zebrafish embryos were maintained in E3 embryo medium (5 mM NaCl, 0.17 mM KCl, 0.33 mM CaCl_2_, 0.33 mM MgSO_4_) in 10 cm Petri dishes at 28 °C. The *tp53* knock-in mutant fish were generated in the *casper* strain^20^ and the *cdh5 G767S* knock-in line also contained *fli1a:EGFF*^21^ transgene.

### Design of sgRNAs and mutation donor ssODNs

The process of sgRNA design for the purpose of replacing particular nucleotides in the endogenous genes (point mutation knock-in or knock-in for short thereafter) involves several bioinformatics analyses. We initially performed protein sequence alignments using NCBI BLAST^22^ of zebrafish proteins to the corresponding human proteins and identified which residues in zebrafish proteins correspond to amino acids mutated in human proteins. Exons containing the amino acid codons to be modified were located in the genomic and cDNA sequences. The sgRNAs were identified using SSC^23^ for efficiency prediction and CC-Top^24^ for off-target prediction. sgRNA sites were also mapped to both genomic and cDNA sequences of the genes and placed into the context of amino acid codons. We then introduced the desired codon mutations and inactivating PAM site or spacer site mutations in the sgRNA sites into the gene and cDNA sequences *in silico* for the genes to be modified. Importantly, sgRNA site mutations were synonymous. The *tp53* ssODNs also contain artificial silent mutations to introduce restriction sites BanI and MspI, but for other knock-ins we did not pursue this strategy. For producing the sense ssODNs we copied 123-136 nt from the *in silico* modified genomic sequence centered on the desired mutation and ensured that the shorter homology arm from the cut site was 60 nt. The anti-sense asymmetric oligos were generated by copying 36 nucleotides to the 5’-end from the cut site of the DNA strand non-complementary to the sgRNA and 90 nucleotides from the same strand in the other direction. This sequence was then reverse-complemented and ordered. These oligos were ordered from Integrated DNA Technologies as 4 nanomole Ultramer oligos. The *lmna* R471W oligos were 90-nucleotide sense oligos which contained the codon mutation and sgRNA site mutations with homology arms of 60 and 30 nucleotides. The PS-modified oligo had the two phosphates on each end of the oligo replaced with a phosphorothioate.

### Synthesis of sgRNAs and Cas9 mRNA

The corresponding sgRNAs were generated by performing an overlap-extension PCR as described previously^5^ with the sense sgRNA oligos (Table S1), each coupled with Rev-sgRNA-scaffold oligo using Taq DNA polymerase (ABM, G009) and then using the resulting PCR product for *in vitro* transcription using MEGAshortscript T7 kit (Thermo Fisher Scientific, AM1354). The sgRNA was purified according to the kit instructions. Cas9 mRNA was made from pT3TS-nCas9n plasmid^25^ after its linearization with XbaI using mMessage mMachine T3 kit (Thermo Fisher Scientific, AM1348) and purified with LiCl precipitation according to the kit instructions.

### Knock-in microinjections

All knock-in injections were performed with Cas9 mRNA at 300 ng/μL, sgRNA at 150 ng/μL and single-stranded oligos at 1 μM into 1-cell zebrafish embryos. Assessment of sgRNA efficiencies was performed using either T7 Endonuclease I (NEB, M0302S) digestion according to the manufacturer’s protocol or using heteroduplex mobility assay (HMA) similarly to what was described previously^26^. All of the oligos mentioned in this section are listed in Table S1.

### Preparation of embryonic and adult samples for genotyping

Samples for genotyping of embryos (2-3 dpf) were prepared by treating embryos with 0.02 % Tricaine in fish medium, transferring them into PCR tubes and replacing the fish medium with 40 μL of 50 mM NaOH, heating at 95 °C for 10 minutes, vortexing, cooling and neutralizing with 4 μL of 1 M Tris-HCl pH 8.0 as described previously^27^. The same procedure was used for genotyping adults except that samples taken were fin clips. Extracts from embryos pools were prepared by combining 50 embryos into a single sample and adding 1 ml of 50 mM NaOH preheated to 95 °C and then following the procedure except that Tris-HCl volume was adjusted to 110 μL.

### PCR assays for genotyping and allele-specific PCR assays

The allele-specific PCR assays that we developed for discriminating between wild-type and point mutation knock-ins are based on several principles. First, the wild-type and knock-in detection primers differ by 2 or more nucleotides (either codon replacement or codon mutation and a PAM site mutation or another silent mutation), one of the mismatches being located at the 3’-most position of the allele-specific primers. Second, the annealing temperature (T_anneal_) for the PCR was typically calculated using NEB Tm Calculator online tool (http://tmcalculator.neb.com/#!/) or alternatively, once a sample was positive for knock-in, the optimal temperature was determined empirically using gradient PCR. This is what was done in the case of the *tp53* R217H knock-in AS-PCR because it initially had high background. Third, we used the touch-down PCR method for all AS-PCRs described in this study, which works as follows: 94 °C for 3 min; **10 cycles**: 94 °C for 30 sec, T_anneal_ + 10 (with 1 °C decrease every cycle), 72 °C for 1 min/kb, **25 cycles**: 94 °C for 30 sec, T_anneal_, 72 °C for 1 min/kb. Finally, all AS-PCRs were done with Taq polymerase because the error rates are not a concern in this method and, more importantly, due to the fact that proof-reading polymerases may remove mismatches between the knock-in primers and wild-type genomic DNA leading to false-positive amplification. T_anneal_ for *tp53* R143H, *tp53* R217H, *cdh5* G767S and *lmna* R471W was 55, 58, 51 and 51 °C, respectively.

### Illumina-based sequencing of amplicons from knock-in embryos to quantify mutation rates

PCR products for HiSeq Illumina sequencing were prepared using primers containing all of the relevant priming adapters and indexes according to the relevant experimental design (single or triplicate samples) (Table S1). Q5 High-Fidelity 2x Mastermix (NEB, M0492) was used for amplifying PCR products for these high-throughput sequencing analyses. PCR products from individual biological samples were amplified using different indexed primers and then pooled into sequencing samples. The sequencing and initial data processing were done by the Next Generation Sequencing Facility of The Centre for Applied Genomics in Toronto, ON. FASTQ files containing paired sequencing reads were assembled by FLASH^28^, mapped to the reference amplicons using bowtie2 software^29^ and SAM files were generated. The SAM files were then processed using custom Python scripts (https://github.com/SergeyPry/knock-in_analysis) to categorize the editing events. The counts of different event categories were processed and plotted using R scripts (https://github.com/SergeyPry/knock-in_analysis).

### cDNA cloning and sequencing

The heterozygous fish carrying knock-in mutations were outcrossed and the embryos were grown to 30 hpf. RNA was extracted from 50 embryos using RNeasy Mini kit (QIAGEN, 74104). cDNA was produced by mixing 10 μL of total RNA with 4 μL of 2.5 mM dNTP and 2 μL of 100 μM oligodT(18), heating at 70 °C for 10 min and cooling on ice. We then added 2 μL of M-MuLV buffer (NEB, M0253S), 0.25 μL of Protector RNAse Inhibitor (Roche, 03335399001), 0.25 μL of M-MuLV reverse transcriptase (NEB, M0253S) and 1.6 μL of water. The synthesis reaction was incubated at 42 °C for an hour and for 10 min at 90 °C. We used Q5 Hot Start High-Fidelity 2X Master Mix (NEB, M0494S) for amplifying cDNA fragments for *tp53* using p53cDNA_for and p53cDNA_rev primers (Table S1) and for *cdh5* gene using cdh5_lastExon_for and cdh5_lastExon_rev (Table S1). The PCR protocol used was 98 °C for 30 seconds, 35 cycles of: 98 °C for 10 seconds, 64 °C for 30 seconds, 72 °C for 30 seconds/kb and the final extension at 72 °C for 2 minutes. The whole PCR reaction was gel-extracted using QIAGEN Gel Extraction kit (QIAGEN, 28704). The purified PCR was cloned into pME-TA using a previously published method^30^. The clones were screened by Taq-based colony PCR using universal M13 primers and then sequenced.

## Results

### Enhanced detection of ssODN CRISPR/Cas9 knock-ins in zebrafish using allele-specific PCR

Genetic diseases in humans are frequently caused by point mutations, but until recently these mutations were modeled in laboratory animals using null mutants of the affected genes, which may result in too extreme a phenotype and possibly not recapitulate the true phenotype seen in human patients. Since zebrafish is a prime animal model for understanding human disease by recreating relevant mutations and given the dramatic progress in genome editing, this animal model is poised for development of effective methods for precise point mutant generation. We aimed at creating defined point mutations R143H and R217H in the zebrafish *tp53* gene at the positions equivalent to those most frequently mutated in patients with the Li-Fraumeni cancer predisposition syndrome. In another project, we decided to introduce a specific G767S mutation into *cdh5*, a gene involved in blood vessel development^31^. The design of the targeting strategies is based on the idea of identifying effective sgRNAs binding as close as possible to the codons to be mutated or even overlapping it as in the case of *cdh5* (Fig. 1A). Ultramer ssODNs used for introducing mutations had the length of 126-136 nucleotides and contained desired mutations and PAM site mutations to prevent cleavage by Cas9-sgRNA complexes of the newly mutated genomic sites as well as mutations to introduce restriction sites that could theoretically be used for genotyping (Fig. 1A), with both of these additional mutations silent at the codon level. The two added mutations were used in the ssODNs for *tp53* modification, whereas *cdh5* G775S ssODN contained a single additional silent mutation because the mutation to be introduced was very likely to disrupt sgRNA binding. As a first step for successful defined mutation knock-ins, the activity of sgRNAs on their respective sgRNA target sites was demonstrated using heteroduplex mobility assay based on a shift of heteroduplex DNA upward for *tp53* sgRNAs and using T7 Endonuclease I digestion for *cdh5* sgRNA (Fig. 1B). Several previous studies of ssODN-based knock-ins of defined point mutations in zebrafish used restriction enzymes for detecting and quantifying knock-ins^8,11^, which led us to introduce BanI and MspI restriction sites into *tp53* knock-in HDR donor templates. We performed injections of ssODN alone and also of knock-in mixes for both *tp53* knock-in strategies and then attempted to quantify knock-ins using BanI and MspI restriction enzymes. However, none of the PCR products from uninjected, ssODN-only injected embryos or those injected with knock-in mixes was detectably cleaved by the enzymes having the sites present in ssODNs (Fig. 1C). Although this result may seem surprising, it is likely that most of the knock-in allele copies are in heteroduplexes with the wild-type PCR products and these heteroduplexes cannot be cleaved by the restriction enzymes. With the lack of success of this initial genotyping strategy, we turned to the allele-specific PCR (AS-PCR) technique that has a long record of successful application in genotyping single-nucleotide and other short polymorphisms. AS-PCR requires two sets primers, one primer in both sets matching its binding site perfectly and the other having two versions for detecting either the wild-type allele or a mutant allele. The variant detection primers have their 3’-most nucleotides matching the respective alleles. When AS-PCR is applied to detecting single-nucleotide variants, it is typically necessary to introduce an additional mismatch not further than 3 nucleotides from the 3’ end into both wild-type and variant primers to enable specific detection^18^. However, in our strategy, the multiple closely-spaced mutations in *tp53* or a complete codon replacement in *cdh5* present in the ssODN and the resulting modified genomic regions allow the wild-type and knock-in detection primers to be designed to simply match their respective alleles (Fig. 1D). We applied our AS-PCR strategies to the *tp53* knock-in samples used in Fig. 1C and also to *cdh5* G767S samples. For all 3 knock-in strategies, the wild-type primer sets produced amplicons of the expected size, whereas the knock-in primer sets only amplified correct products in the knock-in samples (Fig. 1E) thus confirming the validity of our approach.

**Figure 1.**
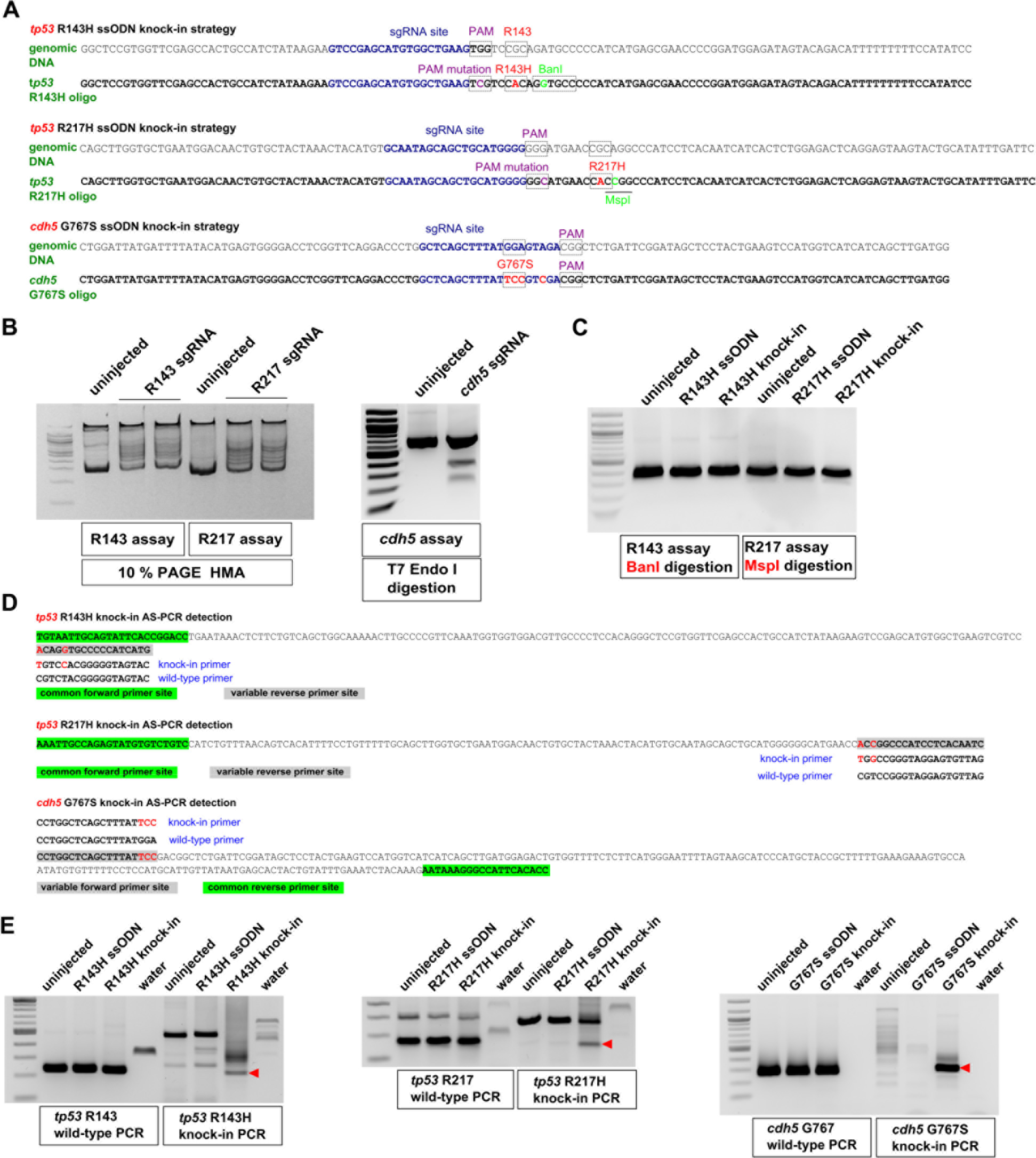
Design of knock-in strategies for *tp53* and *cdh5* point mutations and the AS-PCR assays to detect them. **A.** Genomic sequences and donor oligos are shown for sites of *tp53* R143H, R217H and *cdh5* G767S knock-ins. sgRNA sites are shown in dark-blue, PAM sites and target codons are boxed and their mutations are highlighted in magenta and red, respectively. Mutations leading to introduction of BanI (boxed) and MspI (underlined) restriction sites are highlighted in light-green. **B.** Mutations induced in *tp53* were detected by HMA using 10 % PAGE run. Detection of indel mutations in *cdh5* was performed using T7 Endonuclease I assay. Comparison of PCR product samples from uninjected zebrafish embryos with those injected with respective sgRNAs shows the degree of sgRNA effectiveness. **C.** Restriction analysis of PCR products from uninjected, oligo-injected and *tp53* knock-in injected embryos with enzymes introduced into donor oligos. **D.** Allele-specific PCR assays for detecting introduced knock-ins are based on the idea that when 2 or more nucleotides are different between the wild-type and knock-in alleles, it is possible to design primers that can distinguish the two. Primers common to both wild-type and knock-in assays are highlighted in green and the site for variable primers is highlighted in gray. The wild-type and knock-in primers are indicated below or above the site and the nucleotides corresponding to the knock-in allele are in red both in the amplicon and the knock-in primer. **E.** Example AS-PCRs are shown that were run on the extracts of pooled embryos from uninjected, oligo-injected, knock-in-injected samples and water. Both wild-type and knock-in AS-PCRs are shown, with the wild-type PCRs typically being very strong and indicative of the expected size for the correct knock-in AS-PCR products (indicated with red arrowheads).

### Quantification of knock-in efficiency by next-generation sequencing

Based on the initial results of AS-PCR strategies application, it became very clear that *tp53* knock-ins were significantly less efficient than the *cdh5* knock-in, which prompted us to quantify the exact knock-in efficiency percentages as well as the relative proportions of correct and indel-containing knock-ins. This was done by amplifying PCR products containing Illumina adapters on each side from genomic DNA samples made from embryo pools injected with either *tp53* R143H or R217H knock-in mixes. We obtained 1.86 million reads for *tp53* R143H knock-in and 1.29 million for *tp53* R217H knock-in samples. Since one of the factors affecting knock-in efficiency is the cutting efficiency of Cas9, we quantified the total indel percentages regardless of knock-ins for both R143H and R217H. The total indel percentages for R143H and R217H were 26.1 and 28.4 %, respectively (Fig. 2A). This result shows that the cutting efficiencies by Cas9 in both knock-in experiments were similar. The limitations of current point mutation knock-in approaches are the low percentage of knock-ins and presence of additional undesirable mutations. We therefore divided all of the knock-in events into four classes: correct knock-ins, knock-ins with deletions, knock-ins with insertions and knock-ins in unmapped sequence reads (Fig. 2B, C, D). The unmapped knock-in events likely represent an inappropriate insertion of donor oligos without recombination. The total percentages for R143H and R217H knock-ins were 1.04 and 0.57 %, with the correct knock-ins constituting 83 % of total knock-ins for R143H and 81% for R217H (Fig. 2B). The high relative percentage of correct knock-in reads suggests that more than 80% of recovered alleles with positive genotyping should contain the correctly modified sequences. However, the alleles with indels and especially unmapped reads (Fig. 2C, D) are very variable in sequence as well as size and thus requiring that the potential knock-in animals are both genotyped and sequenced at the knock-in site to verify the correctness of modifications.

**Figure 2.**
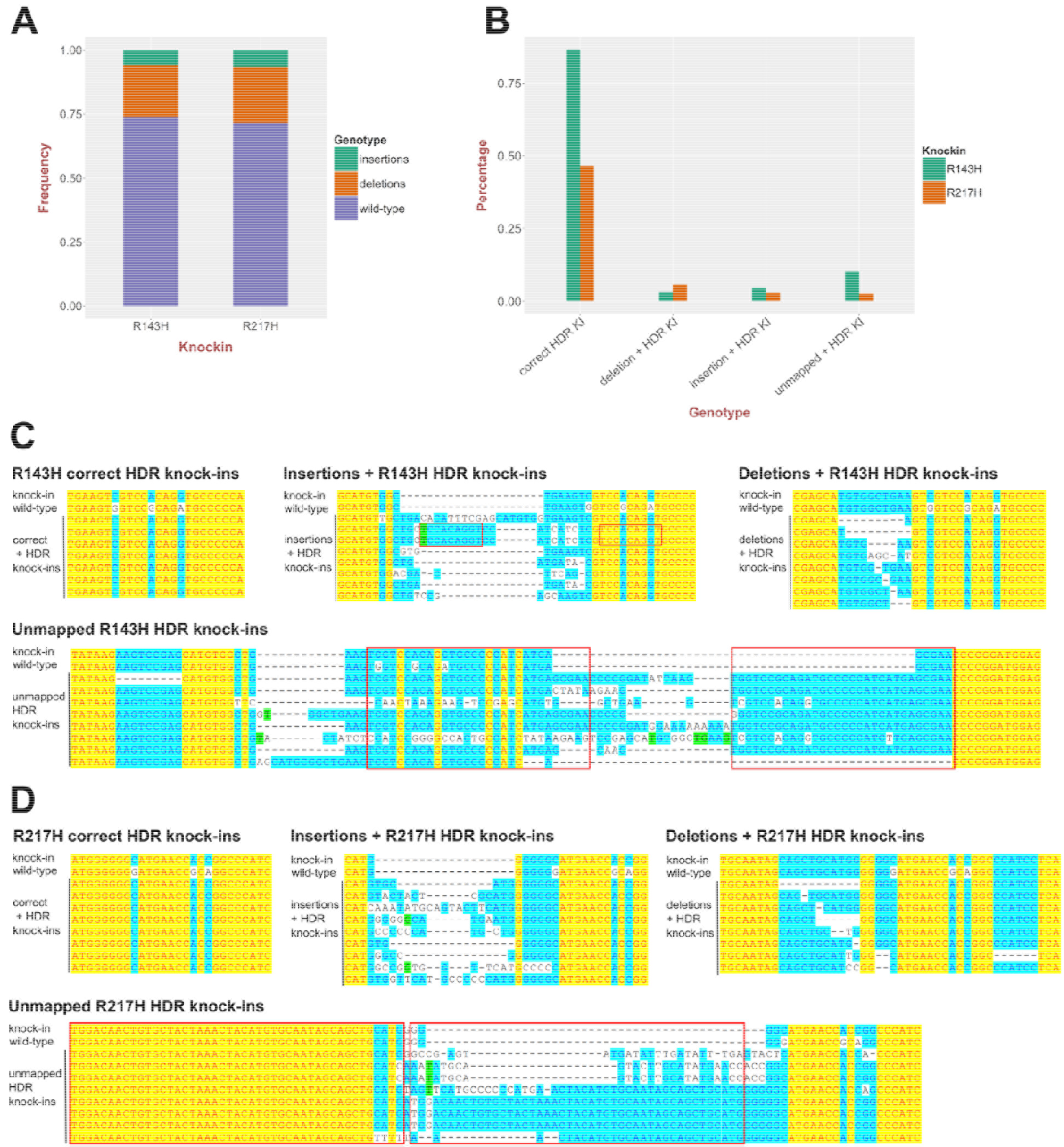
High-throughput sequencing analysis of point mutation knock-ins into the *tp53* gene. **A.** Quantification of the total fractions of small insertions and deletions in *tp53* R143H and R217H knock-in injected embryo samples. The proportion of the reads with deletions or insertions represents a measure of sgRNA activity. **B.** Measurements of R143H and R217H correct homology-directed repair (HDR) knock-ins, knock-ins with additional insertions and deletions as well as knock-in reads with more aberrant complex events (unmapped). **C.** Examples of different classes of R143H knock-ins: correct HDR knock-ins, knock-ins with deletions or insertions and unmapped knock-ins aligned to wild-type and expected knock-in sequences. **D.** Examples of different classes of R217H knock-ins: correct HDR knock-ins, knock-ins with deletions or insertions and unmapped knock-ins aligned to wild-type and expected knock-in sequences.

### Knock-in efficiencies into *tp53* gene are vastly improved using asymmetric anti-sense oligos

Since low efficiency of homology-directed repair can make knock-in generation very laborious and may prevent any positive founder identification as we witnessed in our own initial attempts at *tp53* knock-in generation (Table 1), it is essential to develop approaches to significantly improve the efficiency of ssODN-based knock-ins. Richardson et al., 2016^14^, provided a tremendous advance in this area. Based on extensive biophysical measurements, they found that after Cas9 binds and cuts its genomic DNA site, the DNA strand opposite from the strand binding sgRNA becomes exposed to external molecules. This finding led to the usage of anti-sense asymmetric oligos with the 36-nt homology arm complementary to the strand opposite from the sgRNA-binding site and a 90-nt homology arm corresponding to the sequence on the other side of the Cas9-induced cut. This approach has not been previously employed in zebrafish. We applied this approach to improve *tp53* R143H and R217H knock-ins and compared knock-in efficiencies of the original sense symmetric oligos to those of the new anti-sense asymmetric oligos. After standard knock-in injections with either sense or anti-sense oligos for both R143H and R217H knock-ins, we selected 16 embryos for each type of sample. The semi-quantitative AS-PCR assays show that in cases of both *tp53* R143H (Fig. 3A) and R217H (Fig. 3B), anti-sense oligo knock-ins were dramatically more efficient than the sense ones as reflected in much higher band intensities and fewer bands of larger sizes that very likely correspond to the unmapped type of knock-in reads. To generate quantitative estimates for the improvement in knock-in efficiency by anti-sense asymmetric oligos we generated two sets of samples for R143H and R217H knock-ins with 3 biological replicates for each type of sample (sense or anti-sense). Amplicons with Illumina adapters for these samples were subjected to deep sequencing. For the *tp53* R143H knock-in, overall levels of deletions and insertions were modestly but significantly higher (30.5 % vs 39.2 % for deletions and 7.5 % vs 8.5 % for insertions) for the sense knock-in strategy (Fig. 4A). However, in the case of *tp53* R217H knock-ins the situation was the opposite with both deletions (42 % vs 25.9 %) and insertions (11.8 % vs 5 %) being higher in anti-sense knock-in injections (Fig. 4A). Despite this variation in indel frequency, the anti-sense oligos were significantly more efficient in introducing correct knock-in modifications resulting in 2.72-fold stimulation of *tp53* R143H knock-ins (2 % vs 0.74 %) and 9.5-fold stimulation for *tp53* R217H knock-in (1.92 % vs 0.2 %) (Fig. 4B). The more dramatic improvement of *tp53* R217H may be partially accounted for by the higher cutting efficiencies of Cas9 in those injections. However, differences in cutting efficiencies are not as large as the extent of knock-in stimulation.

**Table 1.** Results of screening point mutation knock-in founders for germline transmission.

| **Knock-in name** | **Oligo used** | **Total number** | **True positives** | **False positives** |
| --- | --- | --- | --- | --- |
| *tp53* R143H | 136-nt sense | 30 | 1 | 2 |
| *tp53* R217H | 136-nt sense | 38 | 0 | 0 |
| *cdh5* G767S | 123-nt sense | 14 | 3 | 1 |
| *tp53* R143H | 126-nt anti-sense | 41 | 2 | 7 |
| *tp53* R217H | 126-nt anti-sense | 22 | 2 | 2 |

**Figure 3.**
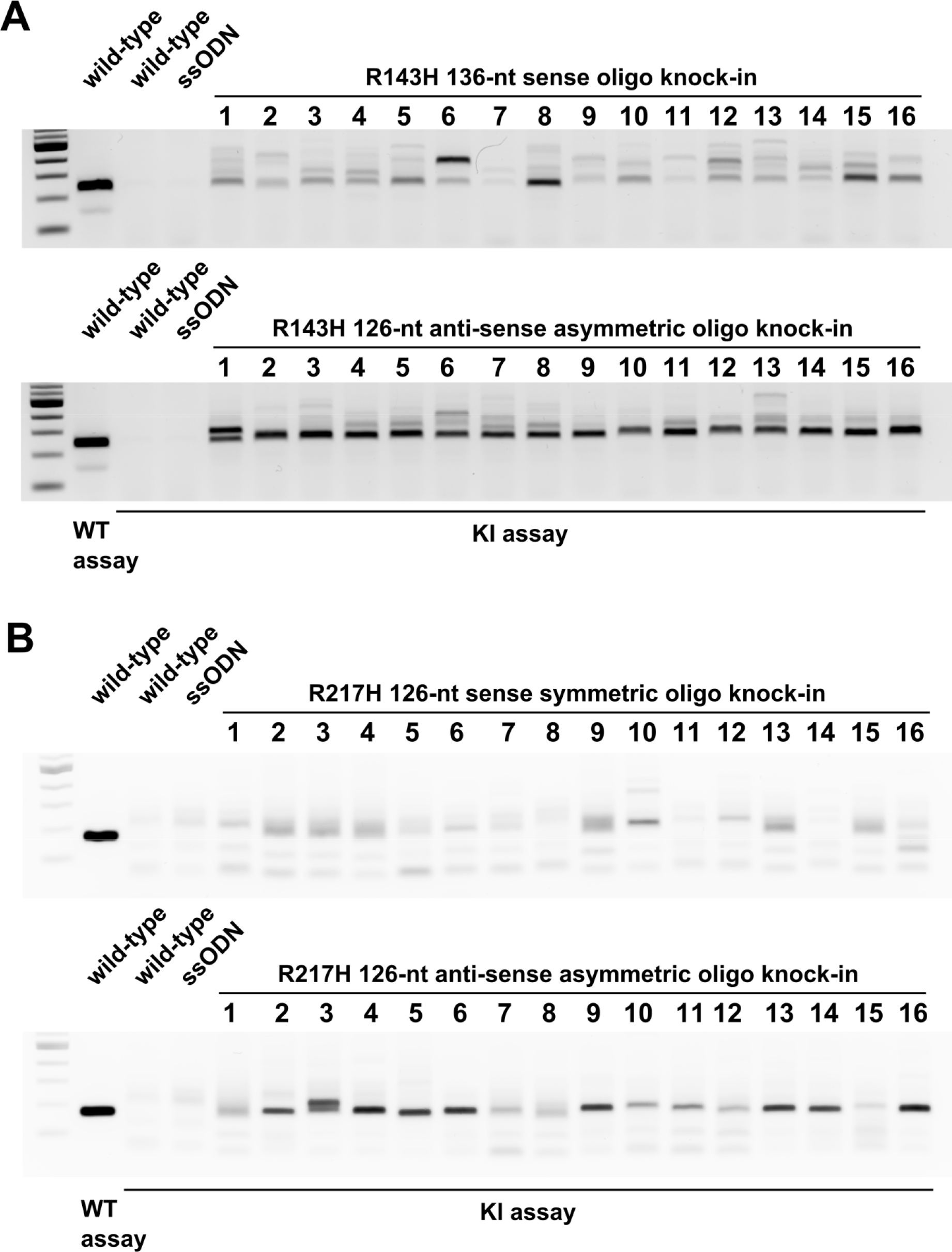
Enhancement of knock-in efficiency by asymmetric oligos anti-sense to the sgRNA sequence. All panels show wild-type embryo genomic DNA extracts run with the *tp53* R143H (**A**) or R217H (**B**) wild-type (WT) assays to serve as controls for PCR product sizes. As negative controls, knock-in (KI) assays for R143H (**A**) and R217H (**B**) were run on wild-type embryos or embryos injected with the respective oligos only. Knock-ins into *tp53* to insert the R143H mutation were performed with either 136-nt sense oligo symmetric around the mutation site or with the 126-nt asymmetric relative to the cut site (36 nt overlapping sgRNA site and 90 nt on the other side) (**A**). In the case of *tp53* R217H, both sense symmetric and anti-sense asymmetric oligos were 126 nt and the asymmetric oligo had the same structure as the asymmetric R143H oligo (**B**). Samples of 16 individual embryos were taken from each knock-in injection and the knock-in assays performed on all of them under the same conditions and at the same time. The gel images shown are representative of at least 3 independent injections and corresponding comparisons of sense and anti-sense oligo knock-in approaches.

**Figure 4.**
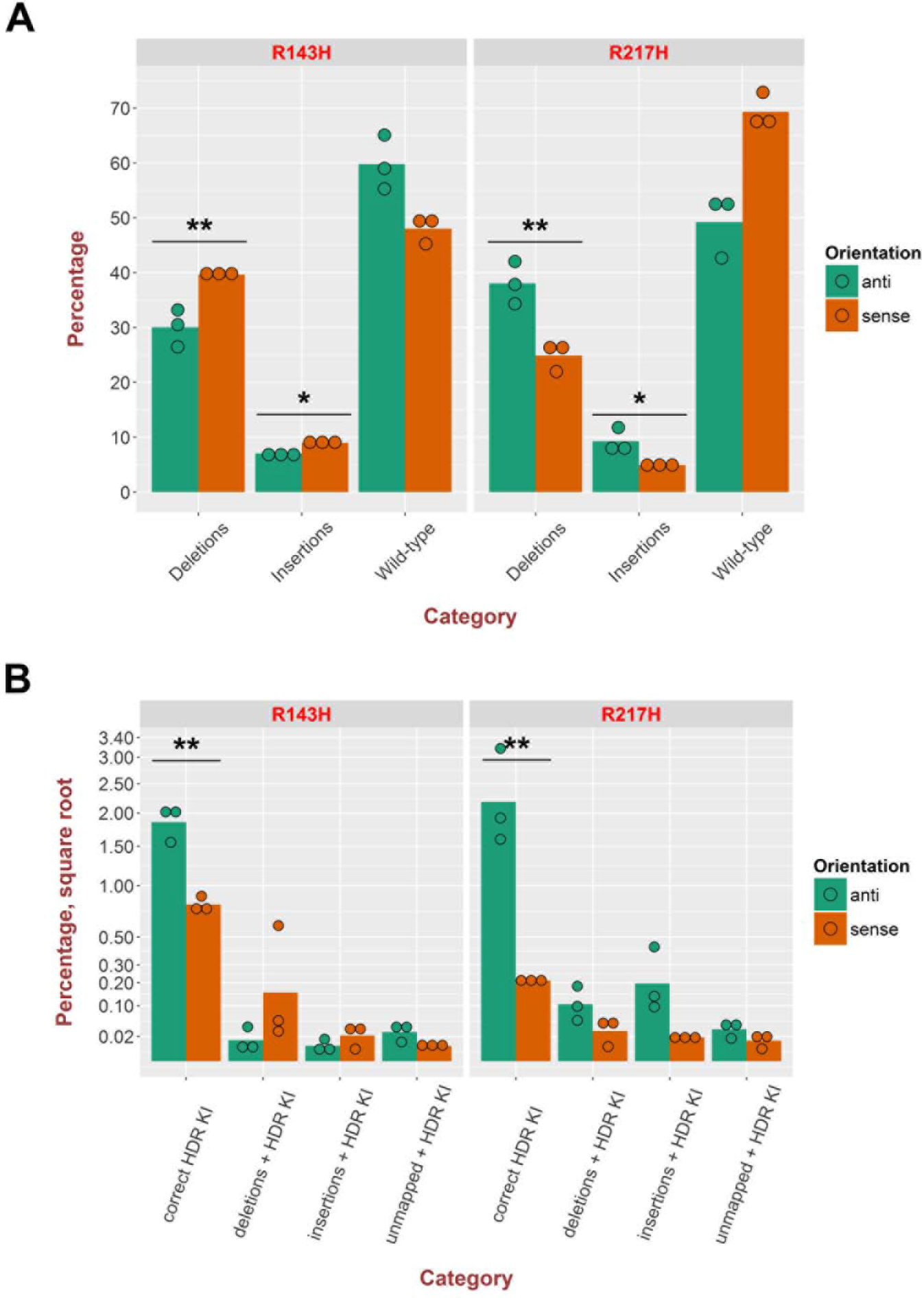
Next-generation sequencing analysis of knock-in efficiency enhancement due to anti-sense asymmetric oligos. **A.** Plot of total percentages of insertions, deletions and wild-type sequence reads at knock-in sites of *tp53* R143H and R217H knock-in samples performed either with anti-sense (anti) or sense ssODNs. **B.** Quantification of percentages of different types of knock-ins at knock-in sites of *tp53* R143H and R217H knock-in samples performed either with anti or sense ssODNs. Square root was applied to the percentage axis for the knock-in plot to better distinguish different values. Bars indicate the mean level for 3 replicates, whose values are plotted with green or orange circles for anti and sense oligos. The significance of differences in insertion, deletion and knock-in rates was determined using t-test and the P-value level is indicated above the respective (** - P-value < 0.01, * - P-value < 0.05).

### A workflow incorporating AS-PCR for rapid and effective isolation of F1 zebrafish carrying correct knock-ins

Although the application of AS-PCR to identify zebrafish (or other organisms) containing correct knock-ins is fairly straightforward, the necessary screening scale due to relatively low modification efficiencies makes the process hard to complete without a defined workflow. In the process of screening more than 100 adult potential founder fish we developed an efficient workflow for isolating correct knock-ins (Fig. 5). The first step of the process is to obtain clutches of F1 embryos from randomly selected potential knock-in founders. Genomic DNA extracts are generated from groups of 50 fish at 24-48 hpf stage from each clutch and the wild-type and knock-in AS-PCR assays are applied to all pooled embryo extracts (Fig. 5A) as has been done for potential *cdh5* G767S (Fig. 5B) and *tp53* R143H (Fig. 5C) knock-in founders. Upon obtaining a positive result for a pooled embryo extract derived from a specific founder, it is recommended to either go back to the clutch derived from this founder or breed the founder again if embryos are not available. One should then prepare 24 individual embryo extracts from a positive founder clutch and at least 2 wild-type embryo extracts. The wild-type embryo extracts are typically run using the wild-type AS-PCR assay to control for the size of PCR products from the knock-in assay and the knock-in assay on wild-type samples ensures that it is specific to knock-in events (Fig 5A, D, E). Example applications of the previous step for *cdh5* G767S and *tp53* R143H positive founders (Fig. 5D, E) show the results that can be expected at this stage of the workflow. To complete the workflow, it is also necessary to amplify the genomic region around the knock-in site (“site assay”) from F1 embryos determined as positive by the knock-in assay. The last step is essential to distinguish between the false-positive and true-positive knock-in embryos and founders. This can be accomplished by assessment of sequencing chromatograms, which show double peaks at the expected positions in the case of true knock-ins (Fig. 5F, G) and wild-type peaks in wild-type samples or false-positive knock-in samples (Fig. 5F, G).

**Figure 5.**
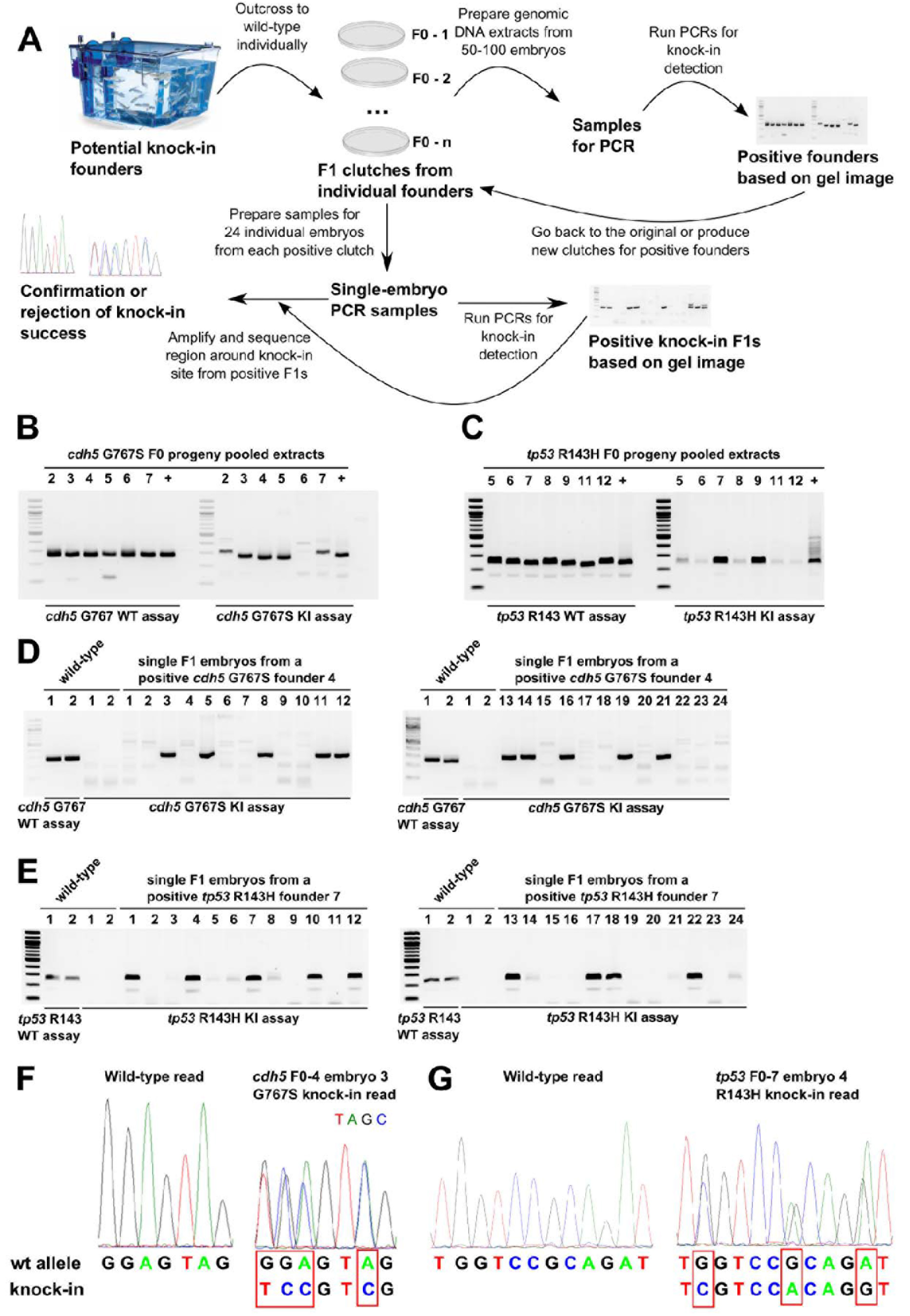
AS-PCR-based strategy enables rapid screening and confirmation of potential founders and knock-in F1 embryos. **A.** In the first step of this workflow, fish injected with the verified knock-in mix (Cas9, sgRNA and oligo) are grown to adulthood and then outcrossed to wild-type fish. The clutches derived from the breedings are used to prepare pooled genomic DNA samples from 50-100 embryos. Wild-type and knock-in PCR assays are then run on these samples to identify the founders with detectable levels of germ-line transmission and to provide the size marker for knock-in AS-PCR products, which should be the same size as wild-type assay products. The information from the first screening round is then used to determine which founders should be bred or which available clutches should be chosen for subsequent screening of 24 individual embryos from each clutch. Knock-in assays are then performed on single-embryo samples and upon obtaining the results, positive embryo extracts are used to amplify a region of DNA surrounding the knock-in site and the resulting PCR products are sent for sequencing to determine if correct knock-in has actually happened. **B, C.** Wild-type and knock-in AS-PCR are shown for *cdh5* G767S (**B**) and *tp53* R143H (**C**) knock-in screening of extracts from embryo clutches produced by potential founders. **D, E.** Screening examples of 24 individual embryos from *cdh5* G767S and *tp53* R143H knock-in founders by AS-PCR. Wild-type and knock-in AS-PCR on two wild-type embryo samples are shown as controls for PCR product size and as negative controls, respectively. **F.** Sequencing chromatograms from wild-type and heterozygous *cdh5* G767S F1 embryos and alignment of the corresponding sequencing reads confirm successful knock-in at the genomic level. **G.** Sequencing chromatograms from wild-type and heterozygous tp53 R143H F1 embryos and alignment of the corresponding sequencing reads confirm successful knock-in at the genomic level.

Although sequencing of PCRs from modified genomic regions is highly suggestive that the correct modification was introduced, it does not constitute proof since it is conceivable that the mutations may interfere with splicing or another process involved in mRNA biogenesis. To prove that the knock-ins we generated correctly modify corresponding mRNAs, we cloned and sequenced cDNA fragments from all three knock-in zebrafish lines. We confirmed that all types of the knock-in cDNA clones are present at the expected frequencies, sequenced 4 wild-type and 4 knock-in clones for each of the knock-ins and aligned the sequences to the theoretical wild-type and knock-in cDNAs as well as to the target and also to flanking exons in the case of *tp53* knock-ins (Fig. 6). The resulting alignments for *tp53* R143H (Fig. 6A), *tp53* R217H (Fig. 6B) and *cdh5* G767S (Fig. 6C) confirm that the mutations were faithfully transmitted to mRNAs.

**Figure 6.**
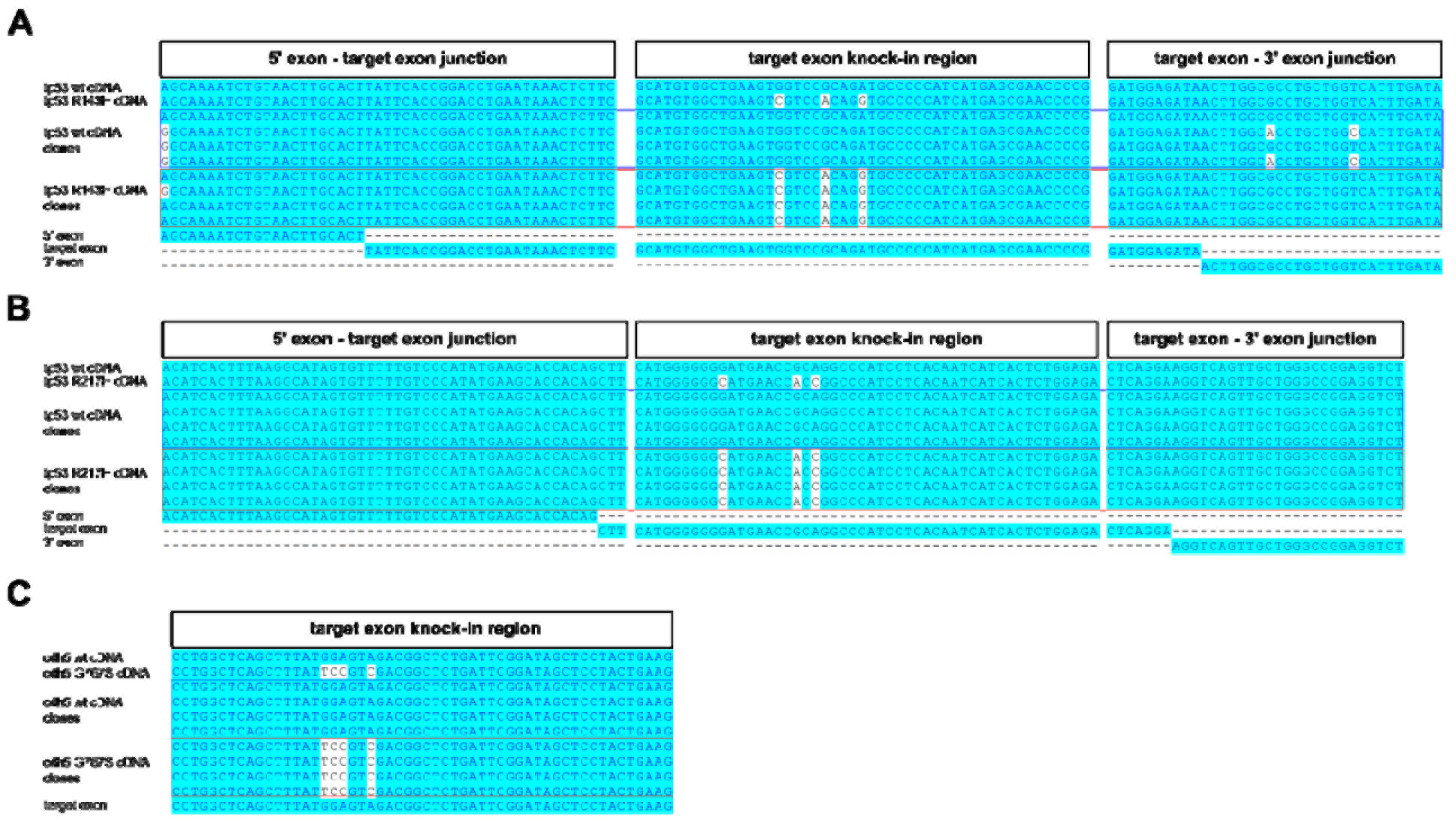
Knock-in analysis in cDNA isolated from F1 carrier zebrafish. Analysis of cDNA cloned from F1 knock-in zebrafish with introduced *tp53* R143H (**A**), *tp53* R217H (**B**) or *cdh5* G767S (**C**) mutations was performed by cloning cDNA fragments, identifying bacterial clones carrying either wild-type cDNA plasmids or those with knock-in mutations introduced, sequencing them and aligning the reads to the expected wild-type and knock-in cDNA sequences. Each of the alignments was performed with wild-type and mutant cDNA reference sequences, 4 wild-type cDNA clones, 4 knock-in cDNA clones and relevant exons. For the *tp53* R143H and R217H knock-ins (**A**, **B**) we show the junction of the 5’ exon and target exon, knock-in region in the target exon and the junction between the target exon and 3’ exon. For the *cdh5* G767S knock-in, we show the target exon alignment because the mutation is at the end of the last exon and therefore unlikely to affect splicing

### False-positive knock-in identification and its possible causes

One common problem affecting many different PCR assays is generation of false-positive results. Contamination with previously generated PCR products or other DNA can be a problem, but it can be quickly resolved by proper procedures, bleach-based cleaning and filter tips. Another more serious problem is when false-positives are generated due to a genetic modification that is different from the intended one. We observed multiple instances of false-positives for AS-PCR when a founder and its F1 embryos were identified as positives by AS-PCR but sequencing the site assay PCR products from positive heterozygous embryos failed to reveal any knock-in events. This occurred for 2 out of 3 founders from the sense *tp53* R143H knock-in and for 1 out of 4 founders of *cdh5* G767S knock-in (Table 1). The original sense *tp53* R217H knock-in strategy failed to produce any positive founders among 38 fish screened, but the improved anti-sense strategy resulted in 2 true and 2 false-positive knock-in founders among 22 fish screened (Table 1). The anti-sense *tp53* R143H knock-in strategy was more efficient than the sense strategy but also produced more false-positives since there were 2 true and 7 false-positive knock-in founders among 41 potential founders (Table 1). Overall, the cumulative founder screening results support the idea that the anti-sense knock-in strategies are more efficient at least for *tp53* gene sites, but also raise concerns about the fact that the false-positive founders account for 25 to 78 % of all positive founders.

Given the potential for confusion and misleading results associated with false-positive knock-ins, we sought an efficient strategy to distinguish between true and false-positive founders. Since the *tp53* R143H strategy included a BanI restriction site that can likely be more tightly associated with the target genomic region, we selected one true knock-in founder (#7) and a false-positive one (#5) for testing (Fig. 7). Some embryos from both founders were expectedly positive for knock-in assay PCR and the true knock-in embryos were indeed less strongly positive (Fig. 7A) than the false-positive ones (Fig. 7B). We then analyzed *tp53* R143 site assays for both sets of embryos before digestion and after digestion by BanI, which revealed that BanI could only digest site assay PCRs of true knock-in embryos (Fig. 7C) but not of false-positive knock-in embryos (Fig. 7D). These results were further confirmed by sequencing true (Fig. 7E) and false-positive knock-in site assay PCRs (Fig. 7F). One possible interpretation of the causes for these problems is that homology-dependent repair (HDR)-independent integration of oligo molecules occurs at other sites in the genome such as at off-target sites, the result of which we propose to call “*trans* knock-in” (Fig. 7G). Eventually, when the PCR is run, single-strand fragments initiated by the knock-in allele-specific primer may pair up with the complementary strand from the endogenous locus and then be extended to produce a regular AS-PCR product (Fig. 7H). Although this model is speculative, it highlights the necessity to perform confirmatory screening and/or sequencing of the knock-in sites after initial AS-PCR hits.

**Figure 7.**
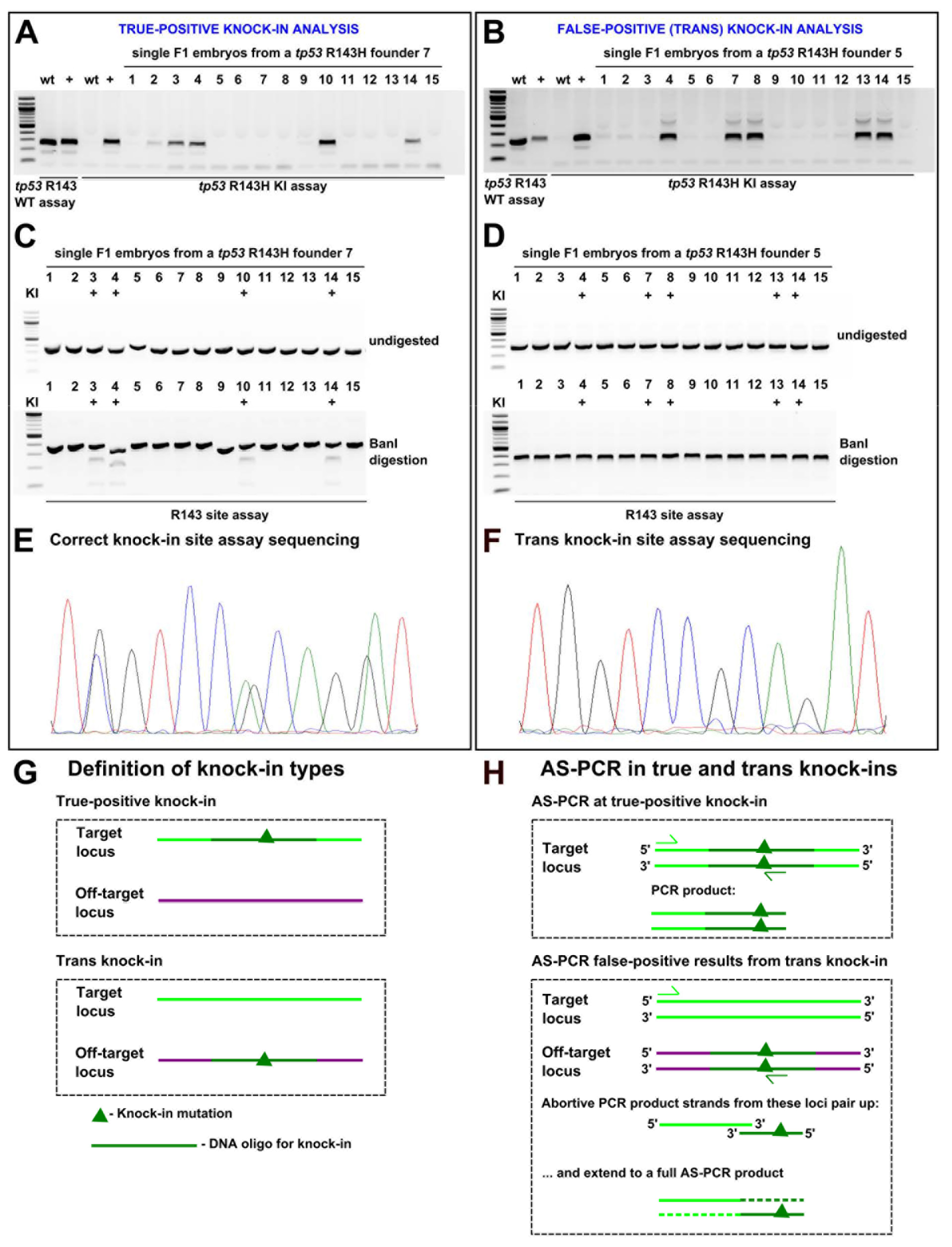
Analysis of true- and off-target (*trans*) knock-ins at *tp53* R143H site. Screening and sequencing verification of true-positive (**A, C, E**) and *trans* (**B, D, F**) knock-ins. **A, B.** A set of 15 F1 embryos from positive founders were analysed using *tp53* R143H knock-in AS-PCR. Wild-type and positive control samples were run with both the wild-type and knock-in PCRs as controls for the size of the PCR product and specificity of the assay. **C, D.** Knock-in site assay PCRs were run on samples from both founders and then either run undigested (upper panel) or digested with BanI enzyme (lower panel) to detect the BanI site expected to be introduced by correct *tp53* R143H mutation knock-in. The samples previously identified as positive for knock-in are marked with “+”. **E, F.** R143H site assay sequencing for true-positive and *trans* knock-in samples. Chromatograms show that in the true-positive knock-in base positions, there are double peaks (**E**), which are absent from the comparable *trans* knock-in read (**F**). **G.** Definitions of knock-in types. In true-positive knock-ins, the targeting oligo modifies the intended locus without off-target insertions, whereas in the *trans* knock-ins, insertion of the oligo occurs at an off-target locus. **H.** A model of how AS-PCR can produce PCR products in both true-positive and *trans* knock-in situations. In the case of true-positive knock-in case, standard PCR mechanism successfully amplifies the expected PCR product. A possible mechanism in the *trans* knock-in case is presented here involving abortive PCR product strands from the wild-type intended knock-in locus and the *trans* knock-in off-target locus. Since the oligo and wild-type locus share significant homology, it is conceivable that very short abortive PCR products from the two loci can pair up and become amplified to the full PCR product in the next cycle thus initiating the exponential cycle of amplification leading to large amounts of PCR product visible as an apparent knock-in band.

### Stimulation of knock-in efficiency and consistency of *lmna* R471W by phosphorothioate oligos

Another attractive, simple and inexpensive way to improve knock-in efficiency is to introduce phosphorothioate (PS) linkages at oligo ends to block exonuclease activity upon them. Based on the observations in cell culture that sense asymmetric and anti-sense asymmetric oligo of 97-nt length were equally effective^12^ for the *lmna* R471W knock-in, we designed 90-nt sense asymmetric oligo versions with or without PS modifications that according to the model suggested by this study will stimulate HDR after a CRISPR-induced double-strand break (DSB) (Fig. 8A, B). Another reason for choosing a sense asymmetric oligo was to uncouple the PS-mediated effects from the stimulation and potential off-target binding of anti-sense asymmetric oligos since the sense oligos do not have complementarity regions for off-target sgRNA sites. We performed a total of 5 *lmna* R471W knock-in experiments with regular (PO) and PS oligos and each time tested 15 injected embryos for both types of injection using knock-in AS-PCR. In all of these experiments, we observed stimulation of knock-ins and a decrease in variation in band intensities after knock-in AS-PCR assay (Fig. 8C). The statistical analysis of measured intensities in 3 of these experiments (44 embryos for each knock-in) showed that there was about 1.4-fold significant increase in average intensity (P-value = 3.9*10^−7^) and a striking shift of measured values in the PS-oligo sample toward the top of the distribution suggesting that in many embryos the stimulation was significantly stronger (Fig. 8). This result shows another way to stimulate knock-in efficiency in zebrafish and the utility of AS-PCR to measure the extent of improvement.

**Figure 8.**
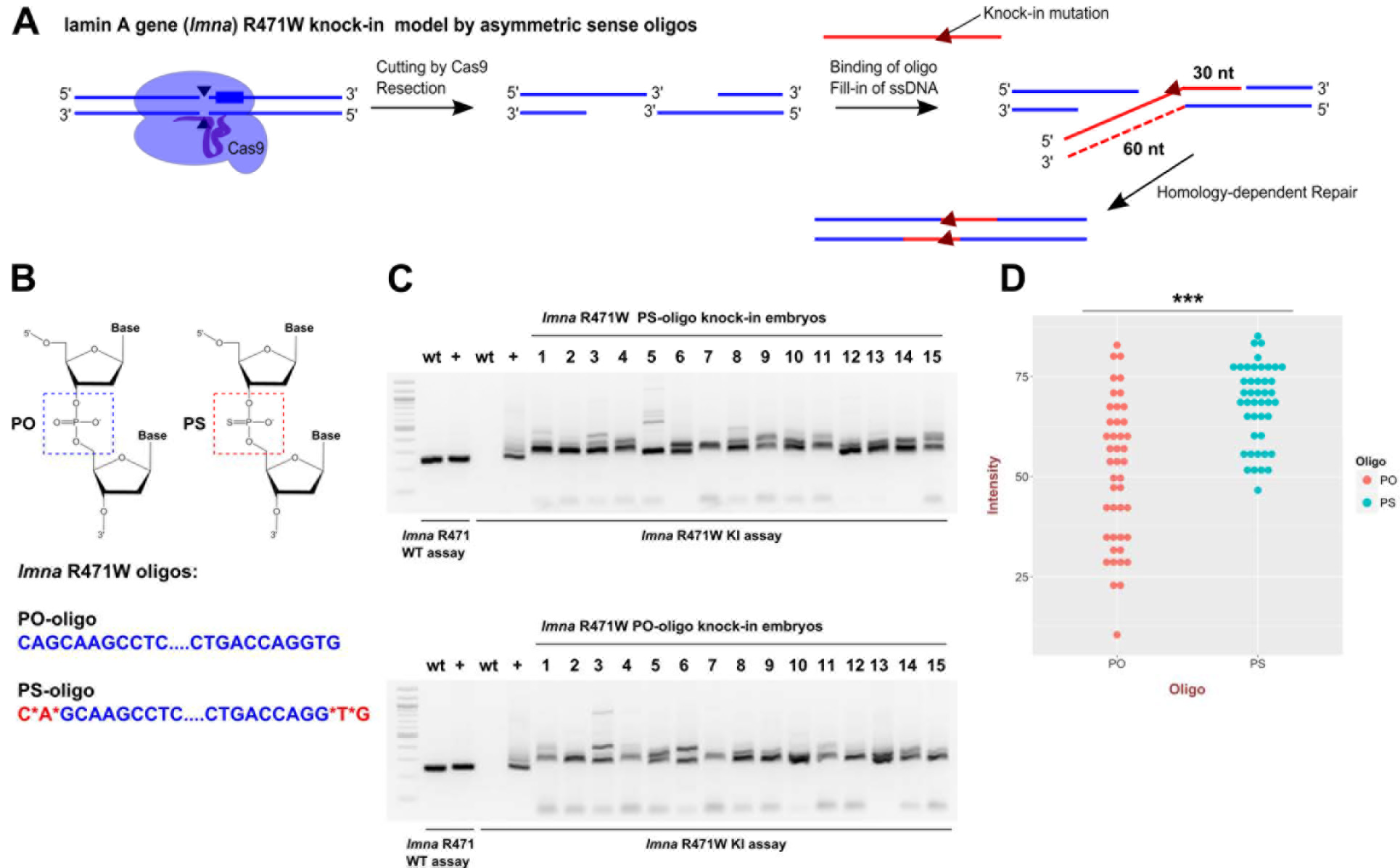
Phosphorothioate-modified oligos improve knock-in consistency and efficiency. **A.** Targeting scheme for introducing R471W knock-in into lamin A/C gene (*lmna*) in zebrafish using an asymmetric anti-sense oligo. Red-colored lines indicate the donor oligo or DNA-derived from it and blue lines indicate genomic DNA. **B.** Chemical structures of the phosphate (PO) and the phosphorothioate (PS) groups show that one of the oxygens in PO is substituted with a sulphur atom in PS. The PS groups were added in the last two chemical bonds on either end of the PS-oligo for *lmna* R471W knock-in and the PO-oligo was synthesized in a standard way. **C.** An example of gel data for AS-PCR analysis of *lmna* R471W knock-ins using PS and PO oligos. WT assay serves as a control for size of the products and the knock-in assay detects the modification. **D.** Graph of measured intensities of AS-PCR signals for 44 embryos for each of the PO- and PS-oligo knock-in injected groups derived from 3 independent experiments. The data are aggregated because there was little variation between experiments. The type of oligo is indicated by color and with x-axis label. The ‘***’ indicate the P-value in t-test of < 0.001 (3.9e-07).

## Discussion

Generation of point mutants using single-stranded DNA oligos and CRISPR/Cas9 genome editing reagents is an emerging technology in zebrafish and other animal model systems. The use of this technique enables single-nucleotide precision of genome editing experiments and will allow generation of specific disease models and precise mutational analysis of biological processes. Despite their obvious promise, point mutation knock-ins remain inefficient in zebrafish and the methods for testing their efficiency remain laborious or not easily accessible to many labs such as sequencing individual plasmid clones by Sanger sequencing and next-generation sequencing of PCR amplicons^11,32^. In an early TALEN-based knock-in study in zebrafish, restriction sites were introduced into specific genomic sites and shown to be digested by the corresponding enzyme^7,8^, but this approach has not yet been shown to work for CRISPR-based knock-ins in zebrafish. The restriction site introduction as a means of genotyping knock-ins is potentially attractive but they will have to be silent mutations in protein coding genes rather than insertions of complete sites and PCR products with silent mutations may behave differently than those with added sequences. In the knock-in projects we describe here, we introduced missense mutations into *tp53*, *cdh5* and *lmna* genes as well as synonymous mutations in PAM or sgRNA sites to prevent Cas9-mediated cutting or to introduce restriction sites for *tp53* knock-ins. Restriction enzymes initially failed to genotype knock-in injected embryos, but were successful at genotyping of F1 heterozygous knock-in embryos. This discrepancy can be explained by the fact that in late cycles of PCR, the strands of different PCR products can be randomly shuffled, from which follows that the fraction of PCR products having both strands containing knock-in mutations has a quadratic dependence on the knock-in allele frequency. Thus, at low (1 and 3 %) knock-in rates (x), only a very small fraction (x^2^) (0.01 and 0.09 %) of total amplicon products can contain fully complementary strands and become digested. By contrast, in knock-in heterozygotes (50% allele frequency), 25% of PCR product can be digested, which was fully consistent with our results. We therefore switched to allele-specific PCR strategy to detect point mutations in all of our knock-ins and have shown that it is very sensitive to knock-in presence at allele frequencies < 0.5 %. Similar detection strategies were previously used for epitope-tagging knock-ins^9,10^, where one of the primers was specific to the tag inserts and the other was outside of the donor oligo region. The epitope tagging detection PCRs and point mutation AS-PCR assays are conceptually similar. However, the relatively small number of nucleotide differences between the wild-type and knock-in alleles can make it hard to avoid background amplification of knock-in assay PCR in wild-type genomic DNA. We employed a touchdown PCR^33^ protocol to make our AS-PCR strategies more specific, which was essential to the success of some AS-PCR assays and improved others.

Overall, our studies focused on point mutation knock-ins revealed three main methods of improving efficiency. The first one was to reduce the distance between the mutation and Cas9 cut site^13^. The first application of AS-PCR also indicated that this strategy may be useful in zebrafish since *tp53* knock-ins where the distances were 10 and 13 nucleotides were much less efficient than the *cdh5* knock-in where the mutation was located exactly at the cut site. This experiment, although suggestive, could be improved by systematic variation of the mutation position relative to the cut site. Another optimization that emerged was the usage of asymmetric oligos. The first such optimization study suggested that only the anti-sense asymmetric oligos can stimulate knock-ins^14^. Knock-in AS-PCRs showed a very strong stimulation using this strategy in the case of *tp53* knock-ins and next generation sequencing confirmed the significance of this result and measured the exact amount of stimulation (ca 3 and 10 fold for R143H and R217H knock-ins, respectively). Another explanation for why asymmetric oligos may function better comes from a well-established model proposing that the protruding single-stranded 3’ regions result from resection of double-strand breaks^34^. The team who explored this model performed knock-ins with multiple 97-nt oligos with different homology arms and they found that the most efficient oligos were 97 nt in length and were designed with the shorter homology arms (30 nt) complementary to the resected single-stranded DNA ends produced after double-strand breaks^12^. These 30-67 asymmetric oligos introduced knock-ins equally well on either side of the DSB thus supporting the resection model much more than the original model proposed by Richardson et al. (2016). Future work will be necessary to establish if stimulation by asymmetric oligos on either strand can be equally effective, but our study provides evidence and an example of how this can be accomplished. In the third optimization we tested phosphorothioate (PS) modification of oligo ends while performinig an *lmna* R471W knock-in. To uncouple potential effects of this modification from those of asymmetric anti-sense oligos and to avoid possible binding of oligos to ssDNA regions at off-target sgRNA sites, we chose asymmetric sense strand oligos of 90 nt with or without two PS bonds at either end of the oligo. Indeed, the PS-modified oligo was significantly more efficient and consistent at introducing knock-ins than the standard DNA oligo. Previously, the group which developed PS-mediated knock-in stimulation could only identify some imprecise knock-in events and did not test for knock-in improvement^15^. We believe that all of these new optimization methods have utility and may even have multiplicative effects when deployed simultaneously. Therefore, at this stage of genome editing technology development, it is advisable to test several versions of oligos incorporating desired optimizations as well as a non-optimized control oligo in order to determine if the optimized versions behave in the expected way.

In the process of genotyping knock-in founders we developed a general workflow to identify true knock-in founders. The unexpected result that emerged from sequencing single F1 embryos was that there were many false-positive or *trans* knock-in founders (25-78 % of total founder number). These could be screened out by sequencing or restriction digests, but they also revealed a weakness of AS-PCR strategies, which can falsely produce a positive signal likely due to hybridization of single DNA strands from abortive PCR products from target and *trans* loci. A possible mechanism of *trans* knock-in origin most likely has to do with off-target sgRNA sites, into which the oligo can ligate. One finding related to this is that the proportion of *trans* knock-in founders was much lower in all of our sense knock-in strategies (3 of 82) than in all of anti-sense knock-ins (9 of 63). A possible explanation is that anti-sense asymmetric oligos will likely have some complementarity to ssDNA regions generated at off-target sites, but this possibility needs further investigation.

In conclusion, we have provided and validated strategies to optimize and enhance point mutation knock-in efficiency in zebrafish. We have demonstrated the benefit of using AS-PCR assays and next-generation sequencing for confirmation and identified the phenomenon of *trans* knock-ins, which can be filtered out using digestions of restriction sites introduced with silent mutations. We envision that this work will make point mutation knock-in generation a straightforward procedure accessible to all zebrafish researchers and beyond.

## Acknowledgements

We would like to thank Gretchen Wagner, Emma Cummings and David Maley for excellent fish care. Sergio Pereira performed Illumina Sequencing of PCR products from knock-in sites.

## Supplementary Material

### Successful optimization of CRISPR/Cas9-mediated defined point mutation knock-in using allele-specific PCR assays in zebrafish

Sergey V. Prykhozhij, Charlotte Fuller, Shelby L. Steele, Chansey J. Veinotte, Babak Razaghi, Johane Robitaille, Christopher McMaster, Adam Shlien, David Malkin and Jason N. Berman

**Table S1.**
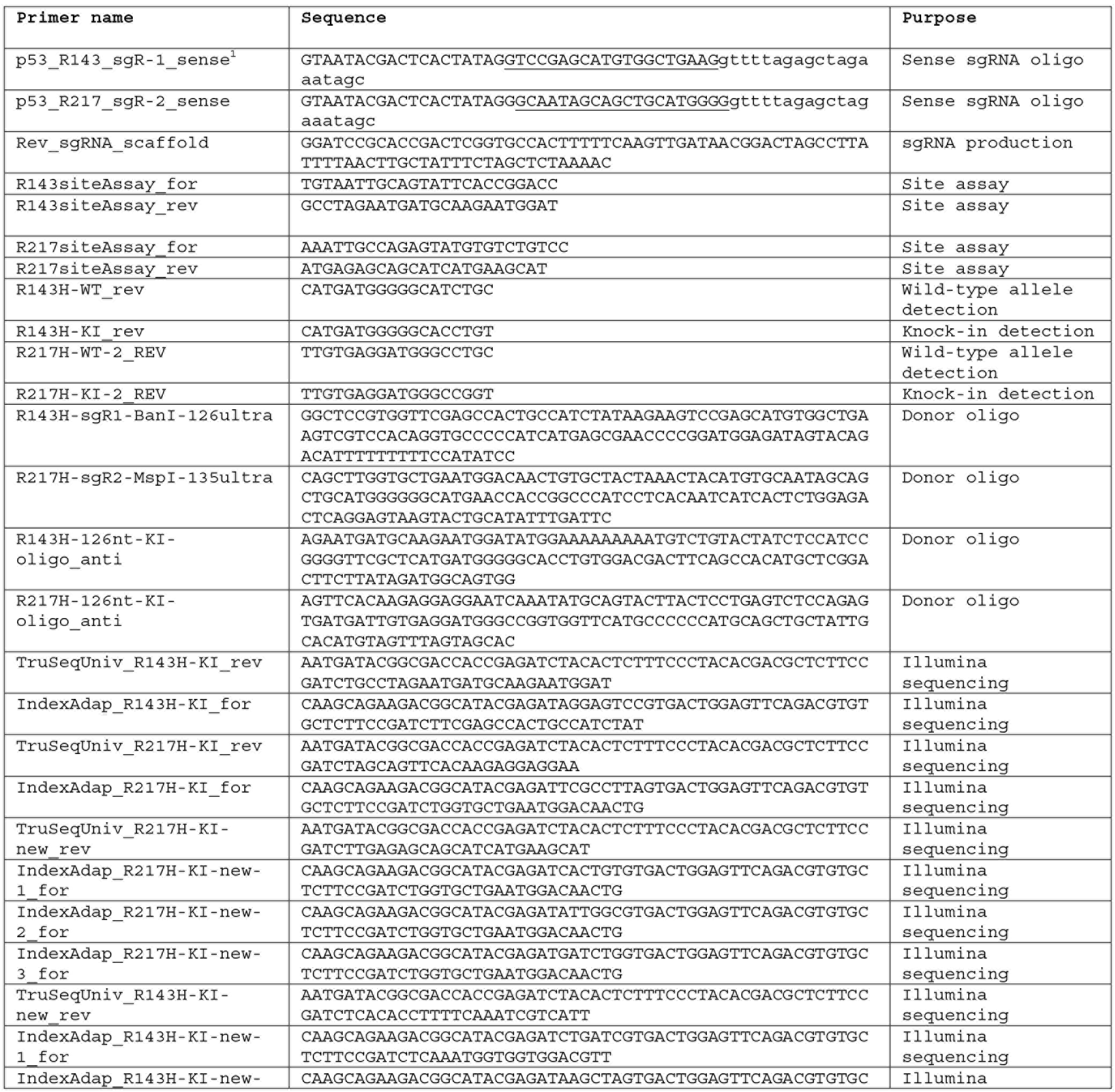
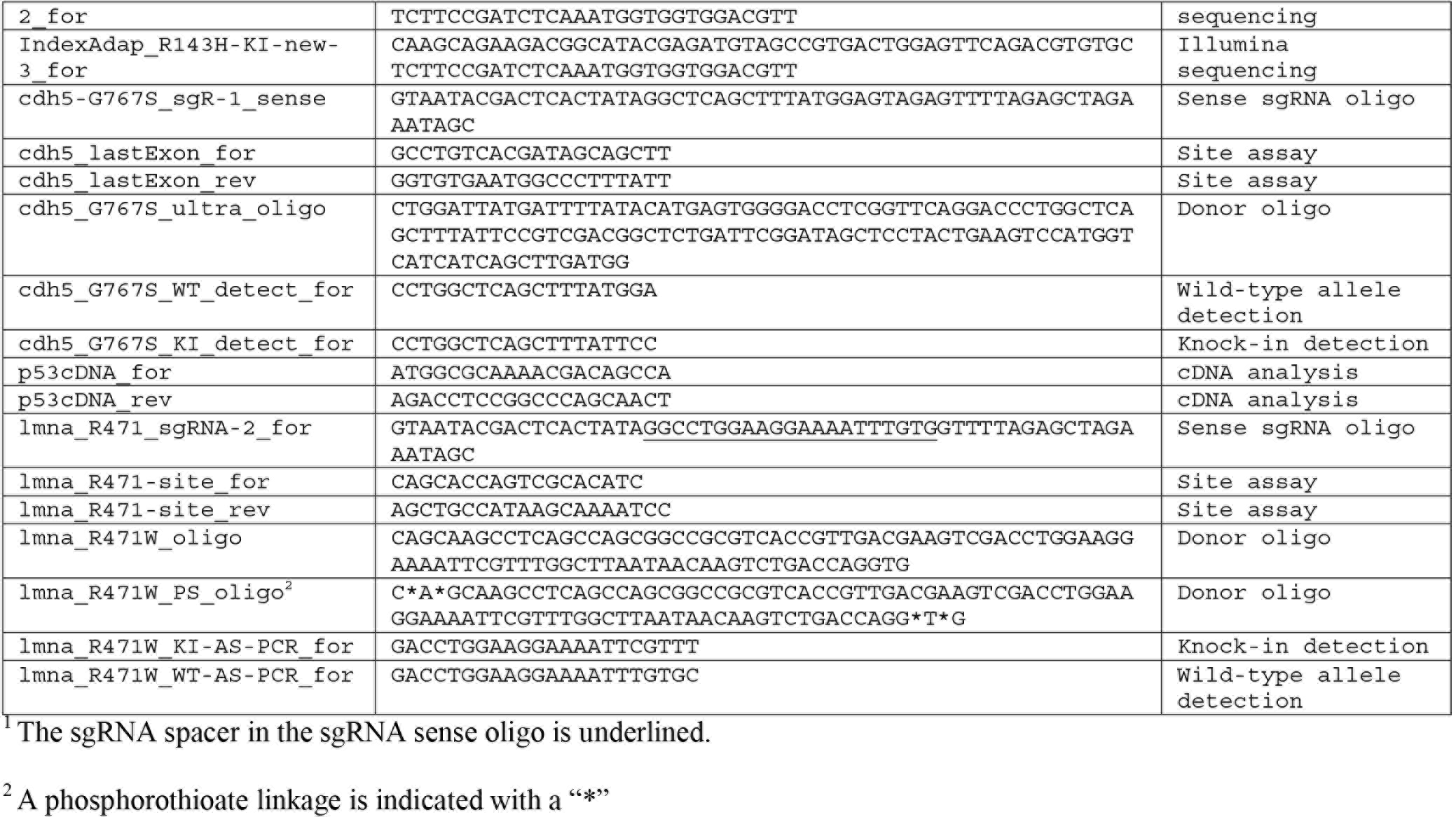
Primers and DNA oligos used in the study.

